# Insights into autophagosome biogenesis from structural and biochemical analyses of the ATG2AWIPI4 complex

**DOI:** 10.1101/180315

**Authors:** Saikat Chowdhury, Chinatsu Otomo, Alexander Leitner, Kazuto Ohashi, Ruedi Aebersold, Gabriel C. Lander, Takanori Otomo

## Abstract

Autophagy is an enigmatic cellular process in which double-membrane compartments, called autophagosomes, form *de novo* adjacent to the endoplasmic reticulum (ER) and package cytoplasmic contents for delivery to lysosomes. Expansion of the precursor membrane phagophore requires autophagy-related 2 (ATG2), which localizes to the phosphatidylinositol-3-phosphate (PI3P)-enriched ER-phagophore junction. We combined single-particle electron microscopy, chemical cross-linking coupled with mass spectrometry, and biochemical analyses to characterize human ATG2A in complex with the PI3P effector WIPI4. ATG2A is a rod-shaped protein that can bridge neighboring vesicles through interactions at each of its tips. WIPI4 binds to one of the tips, enabling the ATG2A-WIPI4 complex to tether a PI3P-containing vesicle to another PI3P-free vesicle. These data suggest that the ATG2A-WIPI4 complex mediates ER-phagophore association and/or tethers vesicles to the ER-phagophore junction, establishing the required organization for phagophore expansion via the transfer of lipid membranes from the ER and/or the vesicles to the phagophore.

## Introduction

Macroautophagy (hereafter autophagy) is a catabolic process essential for the maintenance of nutrition homeostasis and the elimination of cytotoxins, such as damaged organelles, invading bacteria, and aberrant protein aggregates (1, 2). During autophagy, cytoplasmic contents, including cytotoxins, are engulfed in the autophagosome and broken down by lysosomal hydrolases upon autophagosome-lysosome fusion (3). The materials resulting from the degradation, such as amino acids, are recycled. The engulfment of cytoplasmic contents is enabled by the *de novo* formation of the autophagosome. This process involves three dynamic membrane reorganization steps: 1) nucleation of the precursor membrane, called the isolation membrane or phagophore, 2) expansion of the phagophore into a cup-shaped structure, and 3) closure of the open end of the cup-shaped membrane. Despite extensive study, the molecular mechanisms associated with these steps remain elusive due to lack of information regarding the functional roles of the autophagy-related (ATG) proteins (4).

Upon autophagy induction, early ATG factors such as ATG1/ULK kinase, ATG9 membrane protein, and VPS34 PI3 kinase, mediate nucleation of both the early phagophore and the omegasome (5–8). The latter is a PI3-enriched membrane unit observed as an omega/ring-shaped subdomain of the ER by fluorescence microscopy (5) or a cluster of ER-associated thin tubular membranes by electron microscopy (9). Subsequently, the omegasome recruits ATG18/WD-repeat proteins interacting with phosphoinositides (WIPIs) (ATG18 in yeast and WIPI1-4 in mammals) (10–15), the PI3P effector that belongs to the PROPPIN (β-propellers that bind polyphosphoinositides) family (16, 17), and the binding protein ATG2, the largest member of the ATG family. Recruitment of these cofactors leads to the expansion of the early phagophore into a cup-shaped membranous sac. During expansion, the omegasome remains associated with the edge of the open end of the cup-shaped phagophore (5), and ATG18/WIPI and ATG2 also remain concentrated at this ER-phagophore junction (18, 19). When the phagophore has expanded sufficiently, the omegasome starts to shrink and finally disappears as the phagophore is sealed (5). Neither ATG2 nor ATG18/WIPI remain associated with the mature autophagosome (10). The precise spatiotemporal correlation between the localization of ATG2 and ATG18/WIPI and the phagophore/omegasome membrane dynamics suggests that these proteins play direct roles in phagophore expansion at the ER-phagophore interface. Indeed, depletion of ATG2A/B in mammalian cells, which does not affect omegasome formation, leads to an accumulation of small immature phagophores and small autophagosome-like vesicles that are distant from the ER (20).

Although the precise function of ATG2 is unknown, previous studies in yeast suggest that ATG2 may be a peripheral membrane protein (21, 22) that binds directly to membranes (10, 23). In accordance with this described affinity for membranes, mammalian ATG2A/B also localizes to lipid droplets (LDs) and thereby regulate their size (13, 24), as well as localizing at the ER-phagophore junction. These observations collectively suggest that ATG2 may directly mediate a membrane reorganization process, although functional studies have not yielded results to support such claims.

The sequences of ATG2 proteins span ~1600-2300 residues across eukaryotes and contain evolutionarily conserved regions at the N- and C-termini, as well as in the middle of the polypeptide. These domains have been assigned in the Pfam database (25) to the Chorein_N (ID:PF12624), ATG_C (PF09333), and ATG2_CAD (PF13329) families, respectively (Fig. 1A). Chorein_N, and ATG_C share sequence similarity with the N- and C-termini of VPS13 (24, 26), a paralog of VPS13A/Chorein (27). The 200 N-terminal Chorein_N-containing residues and the ATG_C region of ATG2A have been reported to be required for the localization of ATG2A to autophagosome forming sites and LDs, respectively (28). ATG2_CAD contains a highly conserved cysteine-alanine-aspartic acid triad, but its role in autophagosome formation is unknown. In addition, a short region (the residues 1723-1829) preceding ATG_C is also conserved, yet not registered in Pfam. This short region is required for the localization of ATG2A to both phagophores and LDs, and in isolation localizes to LDs (13, 28). We hereafter refer to this region as the C-terminal localization region, or “CLR”. The CLR has been predicted to contain an amphipathic α-helix, indicative of association with membranes (28). Apart from these domains, the regions flanked by Chorein_N and ATG2_CAD, and by ATG2_CAD and ATG2_C in yeast ATG2 were reported to share similarities with the mitochondrial protein FMP27, whose function is unknown, and the Golgi-localized protein of maize APT1, which has been suggested to be involved in membrane trafficking, respectively (23, 29) (Fig. 1A). A fragment containing the APT1 region of yeast ATG2 was shown to interact with membranes containing phosphatidylinositol phosphates, including PI3P (23). However, whether these similarities also apply to higher eukaryotic species is unclear.

**Fig. 1.**
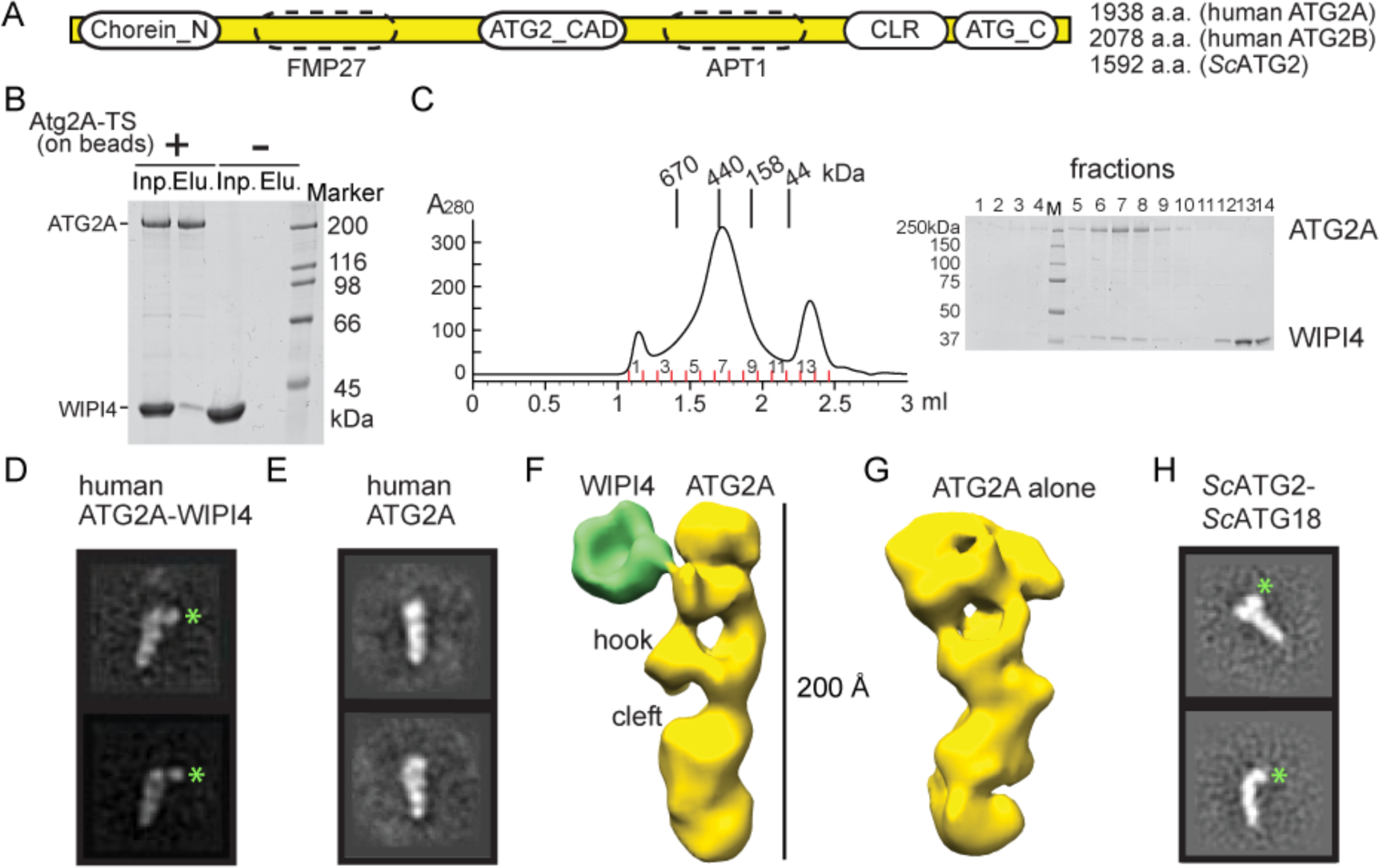
Structural analyses of the human ATG2A-WIPI4 complex and *Sc*ATG2-ATG18 complex by negative stain EM. (A) Diagram of the primary structure of ATG2. The lengths of human ATG2A/B and *Sc*ATG2 are indicated. The regions conserved among all species are indicated by round boxes with solid line. The similarities suggested in *Sc*ATG2 to FMP27 and APT1 proteins are indicated as boxes with dashed line. (B) Affinity capture experiment with ATG2A immobilized on the beads and WIPI4 in solution. (C) Superose 6 size exclusion chromatography profile of the mixture of ATG2A and an excess amount of WIPI4. (D, E) 2D class averages of the ATG2A-WIPI4 complex (D) and ATG2A alone (E). (F, G) Reconstructed 3D structures of the ATG2A-WIPI4 complex (F) and ATG2A alone (G). (H) 2D class averages of the *Sc*ATG2-ATG18 complex. Green symbols in 2D class averages in (D) and (H) indicate the locations of WIPI4 and ATG18, respectively.

In order to better understand the role of ATG2, we characterized human ATG2A in complex with WIPI4 using structural and biochemical methods. Using electron microscopy (EM) and chemical cross-linking coupled with mass spectrometry (CXL-MS), we show that ATG2A has an elongated structure with a WIPI4 binding site at one tip (end). We determined that ATG2A is a bipartite membrane-binding protein that bridges two membranes through interactions at each tip of its elongated structure. Furthermore, we demonstrate that the ATG2A-WIPI4 complex can mediate asymmetric tethering between liposomes with and without PI3P. Placed in the context of the current literature, our findings indicate that the ATG2-WIPI4 complex tethers the PI3P-enriched omegasome to a neighboring membrane(s), such as the ER, phagophore, and other vesicles that may be recruited as a membrane source.

## Results

### Reconstitution and overall structure of the human ATG2A-WIPI4 complex

To enable the structural characterization of ATG2 and investigate its interactions with WIPI/ATG18, we expressed and purified human ATG2A and WIPI4, a pair of proteins that have been reported to strongly interact (12–14), from baculovirus-infected insect cells. The binding was confirmed by an affinity capture experiment in which WIPI4 bound to beads pre-immobilized with ATG2A, but not to beads lacking ATG2A (Fig. 1B). Furthermore, during size exclusion chromatography, the mixture of ATG2A and WIPI4 co-migrated as a single peak (Fig. 1C), thus supporting their ability to form a stable complex. Negative-stain EM studies with the purified ATG2A-WIPI4 complex showed that the particles were monodisperse and homogeneous in size and shape (Fig. S1). 2D class averages of the stained particles revealed that the ATG2A-WIPI4 complex is composed of a rod-shaped protein associated with a small, distinct bead-like feature at one end of the molecule (Fig. 1D). The structural details visible in the 2D averages suggest that the rod-shaped portion of the images corresponds to the multi-domain protein ATG2A. The bead-shaped feature can be provisionally attributed to WIPI4 since the overall shape and size is consistent with its predicted β-propeller fold (30–32). Comparison of these class averages with 2D averages of ATG2A alone supports this proposed organization (Fig. 1E) and establishes that ATG2A and WIPI4 form a 1:1 stoichiometric complex upon reconstitution.

3D reconstructions of the ATG2A-WIPI4 complex and free ATG2A further support the 2D analyses, resolving a rod-shaped ATG2A about ~200 Å in length with a width of ~30 Å. One end of the rod is hook-shaped with a cleft in the middle (Figs.1F and 1G). The WIPI4 density exhibited characteristics that were consistent with a β-propeller, and directly contacts with ATG2A through a thin density (Fig. 1F). This contact likely serves as a hinge through which WIPI4 can adopt a range of orientations relative to ATG2A, as observed in both 2D analyses (SI Movie S1) and 3D reconstructions (Fig. S1). Collectively, these results establish the overall structure of ATG2A in complex with WIPI4. WIPI4 is flexibly associated with ATG2A, inducing no significant conformational change to ATG2A.

### The overall shape is conserved in the yeast ATG2-ATG18 complex

The significance of the interactions between mammalian WIPIs and ATG2A/B has not been thoroughly studied. Much of our knowledge regarding this interaction comes from the studies on the *S. cerevisiae* (*Sc*) ATG2-ATG18 complex. Thus, we investigated whether the structural organization of the human ATG2A-WIPI4 complex described above is conserved in the yeast complex. *Sc*ATG2 is smaller than mammalian ATG2A (Fig. 1A) and appears to bind ATG18 weakly (Fig. S1), which makes the yeast complex more challenging for EM studies than its human counterpart. Nevertheless, we obtained 2D class averages of the ATG2-ATG18 complex (Fig. 1H), which show an elongated object with a bead-like density at one end, very similar to the human complex. These results confirm that the overall structure of the ATG2-ATG18 complex is evolutionarily conserved from yeast to human and indicate that functional studies in yeast are relevant in the context of structural work with the human version.

### Identification of the WIPI4 binding site and insights into the chain topology of ATG2A

We used an integrative approach to gain further structural information on the ATG2A-WIPI4 complex. First, we sought to establish a coarse-grained chain trace of ATG2A and identify the sites of interaction between WIPI4 and ATG2A by CXL-MS. However, there were some foreseeable technical obstacles in performing a CXL-MS analysis of the ATG2-WIPI4 complex. For example, the overall 3D organization of the ATG2A-WIPI4 complex, comprising an elongated ATG2A and the small binding interface between ATG2A and WIPI4, would limit the number of residue pairs that can be cross-linked. Furthermore, the protein complex was prone to aggregation at higher concentrations, limiting the highest protein concentration achievable without introducing aggregation to a moderate level for a CXL-MS analysis. Therefore, to maximize the number of the cross-linked pairs, we performed two cross-linking reactions: a standard amine-coupling reaction with disuccinimidyl suberate (DSS), which cross-links pairs of lysines up to ~30 Å apart (33), and another reaction with the coupling reagent 4-(4,6-dimethoxy-1,3,5-triazin-2-yl)-4-methylmorpholinium chloride (DMTMM) and a linker pimelic acid dihydrazide (PDH), which cross-links pairs of acidic and lysine residues with zero-length (ZL) or two acidic residues via PDH (34). We used relatively low concentrations of the cross-linkers to suppress non-specific intermolecular cross-linking that would cause protein aggregation (see Methods). Despite these technical challenges, mass spectrometry of these samples successfully identified overall twenty cross-linked peptide fragments (Fig. 2A and Table S1).

**Fig. 2.**
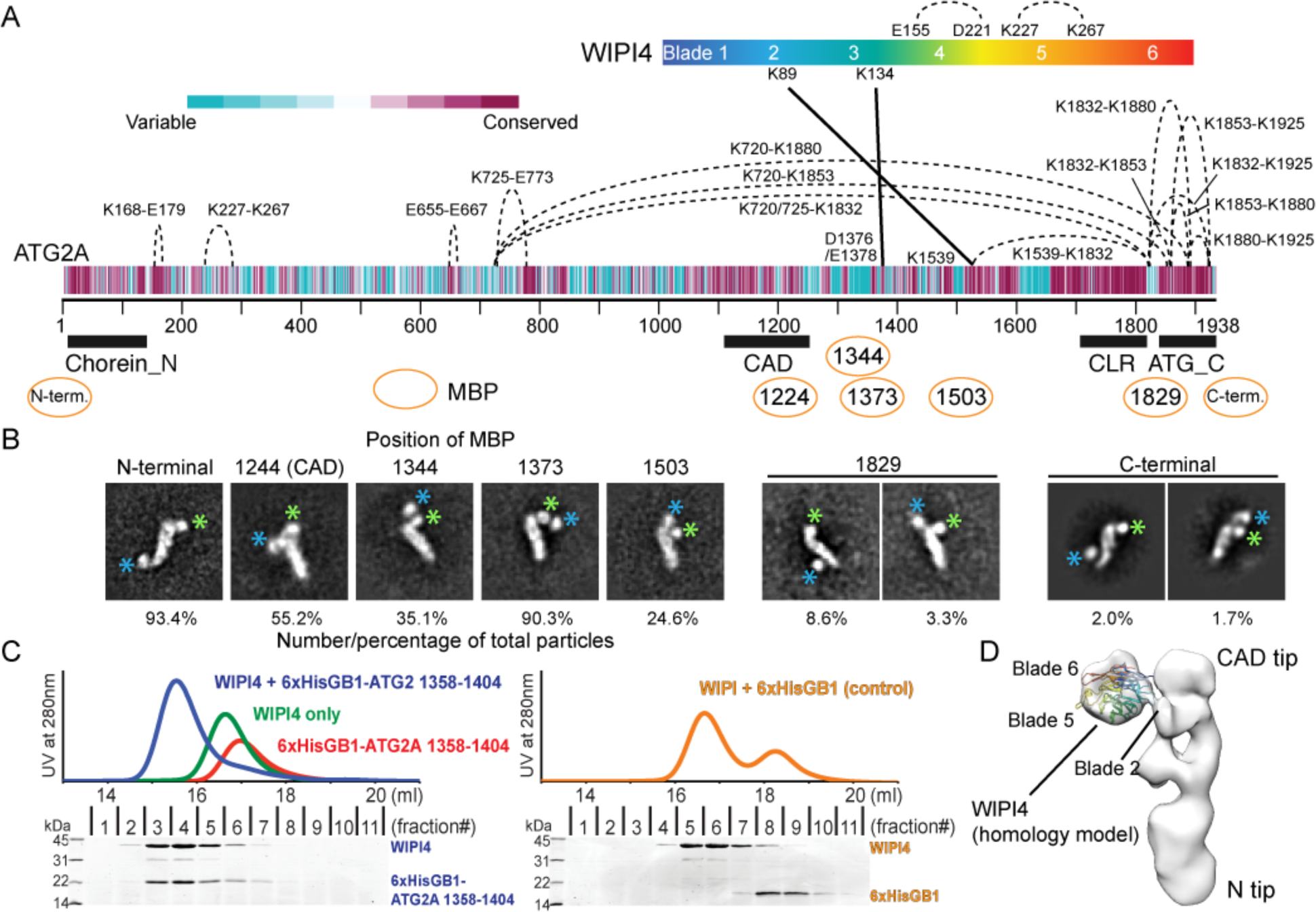
Structural characterizations of the ATG2A-WIPI4 complex. (A) CXL-MS analysis of the ATG2A-WIPI4 complex. Cross-links within ATG2A and between ATG2A and WIPI4 are indicated by dashed and solid lines, respectively. The cross-linked residues are labeled. The diagram of the primary structure of ATG2A is colored per the conservation score of each residue as indicated. The conservation score was calculated on ConSeq server (56). Pfam conserved domains (Chorein_N, ATG2_CAD, and ATG_C) and the CLR are indicated. (B) 2D class averages of MBP-fused or inserted ATG2A in complex with WIPI4. For each construct, a representative 2D class in which the MBP is consistently located adjacent to a tip of ATG2A is shown, with the number of particles (in percentage) that constitute all of the similar 2D classes in each dataset mentioned below. The remaining particles for each dataset where the MBP density is missing in the 2D classes are not shown. The green and blue asterisks indicate the locations of WIPI4 and MBP, respectively. (C) Superdex 200 size exclusion chromatography profiles of 6xHis-GB1-ATG2A (1358–1404), WIPI4, 6xHis-GB1 and their mixtures as indicated. SDS-PAGE on left and right show the size exclusion fraction contents of the sample mixtures containing stoichiometric amounts of WIPI4 and 6xHis-GB1-ATG2A (1358–1404) and of WIPI4 and 6xHis-GB1, respectively. (D) WIPI4-docked electron density map of the ATG2A-WIPI4 complex.

One each of DSS- and PDH-cross-links were identified within WIPI4, and both are consistent with the WIPI4 homology structure model (Fig. S2), validating our experiments. Three cross-links were intermolecular between ATG2A and WIPI4: a DSS-cross-link between Lys1539 of ATG2A and Lys89 on Blade 2 of WIPI4 and two ZL cross-links between Asp1376 or Glu1378 of ATG2A and Lys134 on Blade 3 of WIPI4 (Fig. S2). These data agree with previous work reporting that yeast Atg18 interacts with *Sc*ATG2 through Blade 2 and Loop2, which connects Blades 2 and 3 (30, 35), as well as another study showing that a truncation construct (1–1561) of ATG2A containing the WIPI4-cross-linked residues is able to bind to ATG18 in yeast two-hybrid (36), suggesting evolutionary conservation of this interaction mode. Furthermore, these data also suggest that the WIPI4-bound tip comprises amino acids that are located in a central region of the primary structure. Fifteen cross-links were collected within ATG2A: eleven short-to mid-range (11-93 residues) DSS/PDH/ZL-cross-links, which are indicative of locally-folded subdomains (Fig. 2A), as well as five long-range (293-1160 residues) DSS cross-links between the residues in ATG_C and Lys720/725/1539. SDS-PAGE analysis of the DSS cross-linking reveals an intense band at ~250 kDa (Fig. S3). Because the molecular weight of the ATG2A-WIPI4 complex is ~250 kDa and the ~250 kDa band is the single major band, it seems likely that the five long-range DSS cross-links are intramolecular (Fig. S3), indicating that the folded ATG2A polypeptide adopts a non-linear chain topology. However, a faint smear in the ~500-600 kDa range is also observable, which may result from two cross-linked ATG2A-WIPI4 complexes, raising the possibility that the long-range cross-links could be inter-ATG2A molecules. Attempts to identify cross-links from in-gel digestion of the monomeric band were unsuccessful, probably due to a limited recovery of cross-linked peptides from the gel, preventing us from drawing an unambiguous conclusion.

In order to map the conserved regions of ATG2A as well as the WIPI4-interacting site, a 42 kDa maltose-binding protein (MBP) was fused to ATG2A at the N-terminus, and separately inserted into the ATG2_CAD (after residue 1224) (Fig. 2A). 2D image analyses of negatively stained samples revealed that the MBP fused to the N-terminus localized to the tip opposite the WIPI4-bound tip of the elongated ATG2A (Fig. 2B). The MBP inserted into the ATG2_CAD localized to the same tip that binds WIPI4 (Fig. 2B). These data suggest that the previously described conserved regions are located at opposite ends of the ATG2A rod-like complex, and thus have different functional roles. Hereafter, we refer to the tips of the ATG2A complex as the N tip and the CAD tip. Our identification of the CAD region is consistent with our CXL-MS experiments, as the CAD region is in close proximity to residues that cross-linked with WIPI4. To further confirm this localization, we also inserted MBP into three positions adjacent to the residues cross-linked to WIPI4 (after the residues 1344, 1373, and 1503). As expected, all of these MBP inserts were identified to be in close proximity to WIPI4 in negative stain 2D class averages (Fig. 2B).

In order to identify the C-terminal region, we fused an MBP to the C-terminus and also inserted an MBP between the CLR and ATG_C (after the residue 1829). In contrast to our previous labeling experiments, the MBP molecules in these C-terminal regions were not identifiable in most of the 2D classes. The remaining subset of particles (~12% particles with MBP at the position 1829-1830, and ~4% particles with MBP at the C-terminus (Fig. 2B)) show an additional globular density attached to ATG2A-WIPI4 that we attribute to MBP. However, in both constructs, this MBP density was observed adjacent to both tips of ATG2A-WIPI4 (Fig. 2B). These data do not allow unambiguous localization of the C-terminal regions, but rather suggest that the C-terminus of ATG2A is flexible with respect to the rest of the molecule. In the CXL-MS experiments, three lysine residues in ATG2_C cross-linked to two residues in the middle of the ATG2A sequence (720/725 and 1539) (Fig. 2A). If these cross-links are indeed intramolecular, then this also supports the notion that the C-terminus of ATG2 is flexible and can reach the WIPI4-bound tip.

Given the facts that our EM structural analysis revealed that WIPI4 was flexibly attached to ATG2A, and that β-propellers often bind to peptides, we hypothesized that the WIPI4 binding site is in a flexible linear region of ATG2A. To test this hypothesis, we generated a fragment of ATG2A (residues 1358-1404) containing two of the inter-molecularly cross-linked residues (Asp1375 and Glu1378) as a fusion to the B1 domain of streptococcal protein G (GB1) protein. The GB1-fused peptide co-migrated with WIPI4 in a size exclusion chromatography column and eluted earlier than WIPI4 alone (Fig. 2C). A control using just GB1 and WIPI4 shows no co-migration, demonstrating that this linear region is indeed capable of binding to WIPI4.

Based on these results, we fit a WIPI4 homology model into the 3D EM density, with Blade 2 facing ATG2A (30, 35) and the membrane-binding surface of WIPI4, including the two PI3P recognition sites (Blades 5 and 6), (30–32) on the opposite side of ATG2A (Fig. 2D).

### ATG2A associates with membranes through its tips

Next, we sought to characterize membrane binding by ATG2A by performing a liposome flotation assay using a Nycodenz density gradient (37). Small unilamellar vesicles (SUVs) and large unilamellar vesicles (LUVs) were prepared by sonication and extrusion methods, respectively. These vesicles were mixed with ATG2A in the presence of Nycodenz, floated to the top of a gradient by centrifugation, and subsequently collected and further analyzed. ATG2A rose to the top of the gradient only in the presence of liposomes (Fig. 3A), confirming direct membrane binding. The recovery of ATG2A proteins was substantially higher with SUVs than LUVs (14-32 fold with 1,2-dioleoyl-*sn*-glycero-3-phosphocholine (DOPC) vesicles), suggesting that ATG2A prefers binding to highly curved membranes. Additionally, incorporation of negatively charged lipid (1,2-dioleoyl-*sn-*glycero-3-phospho-L-serine: DOPS) into the liposomes increased ATG2A-liposome association 1.3-9 fold, suggesting some electrostatic contribution to this interaction.

**Fig. 3.**
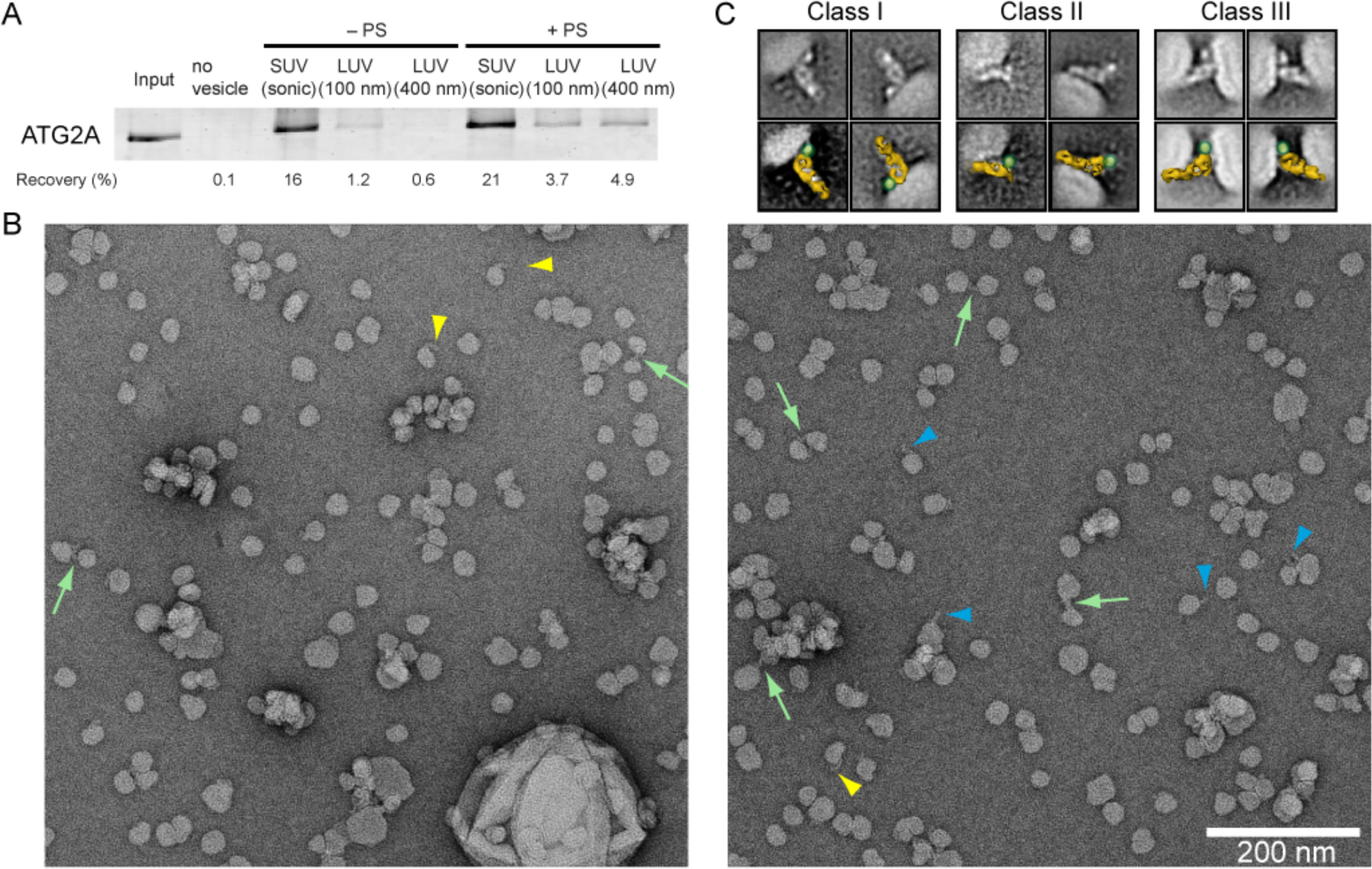
Interaction between ATG2A and liposomes and its visualization by EM. (A) Liposome flotation assay with 50 nM ATG2A. The liposomes composed of 99% DOPC and 1% DiD (indicated as “–PS”) or 74% DOPC, 25% DOPS and 1% DiD (“+PS”) were prepared by sonication (SUVs) or extrusion (LUVs) using 100 or 400 nm filters. The inputs (4%) and the top layers after the centrifugation (24%) were loaded onto SDS-PAGE. The percentage of ATG2A recovered in each of the top fractions was quantified and is shown below the gel image. (B) Micrographs of the negative stained ATG2A-WIPI4-SUV complex. Colored arrowheads and arrows denote an elongated object that emanates perpendicularly (blue arrowheads) or tangentially (yellow arrowheads) from an SUV, or is tethering two SUVs (green arrows). (C) 2D class averages of the ATG2A-WIPI4-SUV complex shown without (top) and with a manually placed 3D model of ATG2A (bottom, shown in yellow) together with a green dot on the WIPI4 density.

To better understand how ATG2A associates with membranes, we performed negative stain single-particle EM analysis on the ATG2A-WIPI4 complex bound to SUVs. Because ATG2A is a thin, somewhat featureless rod, we foresaw difficulties in clearly visualizing such proteins on large membrane surfaces. Therefore, we added WIPI4 to serve as a molecular marker, allowing us to unambiguously determine the orientation of ATG2A bound to the liposomes. We used SUVs composed of DOPC and DOPS, but not PI3P, to avoid any effects introduced by the WIPI4-PI3P interaction. Because SUVs produced by sonication were highly heterogeneous in size, we generated more homogenous SUVs using a dialysis methodology (38). We performed a flotation assay with WIPI4 and these SUVs and confirmed that WIPI4 does not bind to these SUVs (Fig. S4). In the raw micrographs of the ATG2A-WIPI4-SUV complex (Fig. 3B), we observed elongated objects resembling ATG2A associated with either one or two liposomes, as well as clustered liposomes. Focused 2D analyses on the elongated object produced averages containing features consistent with the previously observed ATG2A-WIPI4 complex, including the characteristic hook and cleft (Fig. 3C). The 2D classes could be categorized into three major structural classes of the protein-SUV complexes. In the first class, WIPI4 and the CAD tip of ATG2A are bound to the membrane, with the long axis of ATG2A aligned roughly orthogonal to the membrane, positioning the N tip away from the membrane. In the second class, ATG2A is bound to the membrane through the N tip, with the WIPI4 directed away from the membrane. These data indicate that both tips of ATG2A can interact with membranes independent of each other. Strikingly, particles belonging to the third class can be described as a combination of the first and second classes; ATG2A is bound to one liposome through the CAD tip and to another through the N tip, spacing neighboring liposomes ~10-15 nm apart. These EM data thus suggest that ATG2A is a bipartite membrane-binding protein that can bridge two membranes.

### ATG2A tethers SUVs

To confirm the membrane tethering activity of ATG2A, we examined the effect of the presence of ATG2A on the size distribution of liposomes using dynamic light scattering (DLS). The DLS profile of the SUVs markedly shifted to larger sizes upon incubation with ATG2A (Fig. 4A), whereas those of the LUVs (100 and 400 nm) did not change (Figs.4B and 4C). To confirm that the increase in the SUV particle size was due to liposome clustering mediated by the protein, we added proteinase K to the final sample of the ATG2A-SUV mixture and monitored its effect. Upon incubation, the observed particle size decreased to its original dimensions (Figs. 4D and 4E), demonstrating that homotypic tethering mediated by ATG2A resulted in the clustering.

**Fig. 4.**
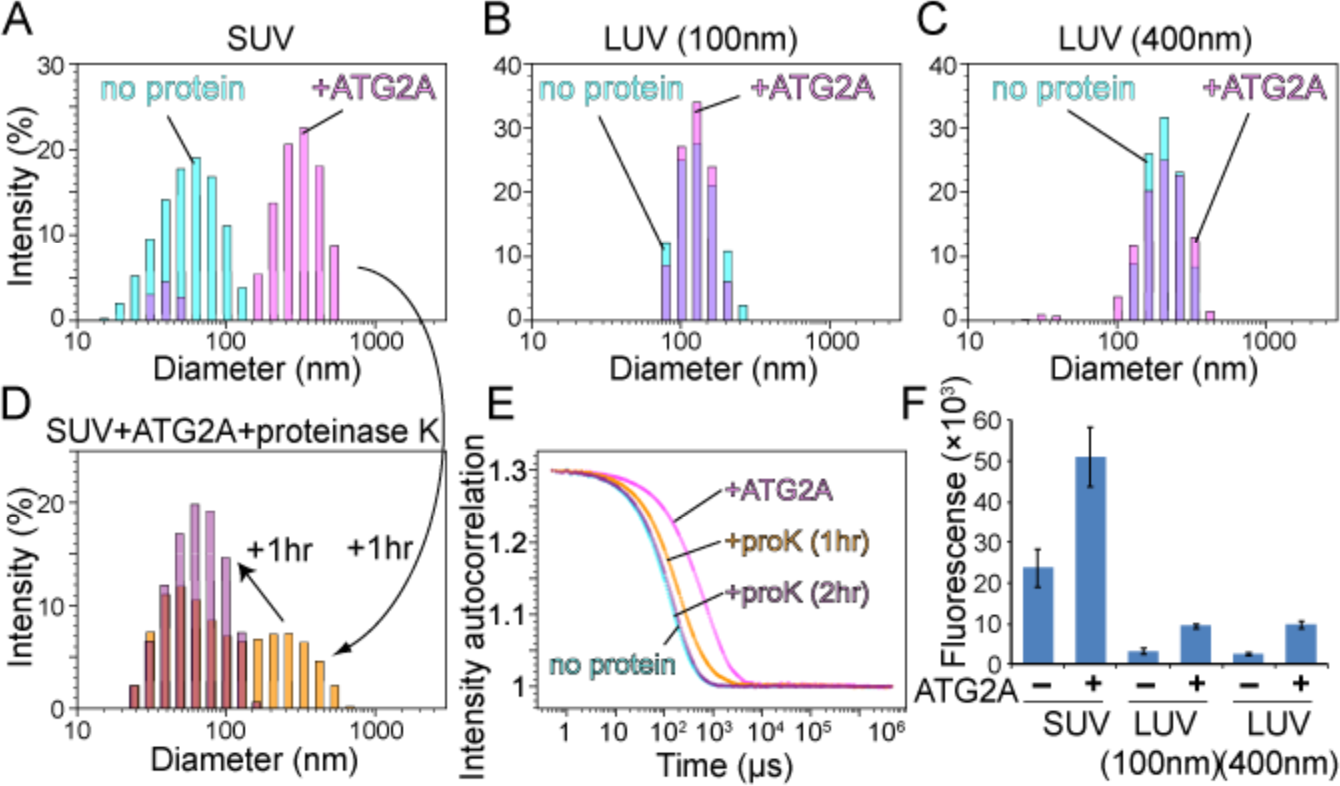
Membrane tethering by ATG2A. (A-C) The DLS profiles of SUVs (A), LUVs (100 nm) (B), and LUVs (400 nm) (C) in the absence (cyan) or presence (magenta) of 200 nM ATG2A. All liposomes consisted of 75% DOPC and 25% DOPS. The samples were incubated for 1 hr before the measurements. (D) The final sample of (A) was mixed with proteinase K and re-measured upon 1 (yellow) and 2hr (purple) incubation. (E) Auto-scaled autocorrelation functions of the four DLS measurements with SUVs (B, D) are plotted. (F) Fluorescence liposome tethering assay. Liposomes consisting of 73.3% DOPC, 25% DOPS 0.2% biotinylated lipids, 1.5% Rhodamine-PE were mixed with liposomes with the same size consisting of 73% DOPC, 25% DOPS, 2% DiD in the presence of 200 nM ATG2A. The fluorescence reports the amount of the DiD-containing liposomes associated with the biotinylated liposomes. For each experiment, the average of three repeats is shown with the standard deviation.

We also performed a fluorescence-based liposome-tethering assay, in which biotin-incorporated liposomes were mixed with liposomes containing fluorescent lipids and separated by streptavidin beads (39). The fluorescence intensity of the beads reports the degree of tethering occurring between these two types of liposomes. The results show that the fluorescence signals of the liposomes, regardless of size, were increased by the presence of ATG2A (Fig. 4F), but the difference between the signals in the presence or absence of ATG2A was ~5 times larger in the SUVs compared to the LUVs. These data suggest that ATG2A is capable of tethering liposomes, with a preference for SUVs. Altogether, the results from flotation assays, DLS, and fluorescence measurements establish that ATG2A can tether small liposomes (i.e., membranes with high curvature).

### The ATG2A-WIPI4 complex tethers PI3P-containing and non-PI3P-containing membranes

We reasoned that in the homotypic tethering experiments described above, ATG2A was unable to tether LUVs (Figs. 4B, 4C and 4F) due to its weak affinity to LUVs (Fig. 3A). Notably, the liposomes used in these experiments lacked PI3P. Because the PI3P effector WIPI4 robustly binds to PI3P-containing LUVs (31), we hypothesized that the ATG2A-WIPI4 complex could also efficiently bind PI3P-containing LUVs, thereby facilitating tethering of these liposomes. To test this hypothesis, we used DLS. As shown in Fig. 5A and 5B, neither WIPI4 nor ATG2A individually changed the size distribution of the LUVs. Thus, the presence of PI3P does not appear to increase the affinity between ATG2A and LUVs. In contrast, the LUV particle size increased markedly in the presence of both proteins (Fig. 5C and 5D), demonstrating that the WIPI4-PI3P interaction triggers the clustering of PI3P-containing LUVs.

**Fig. 5.**
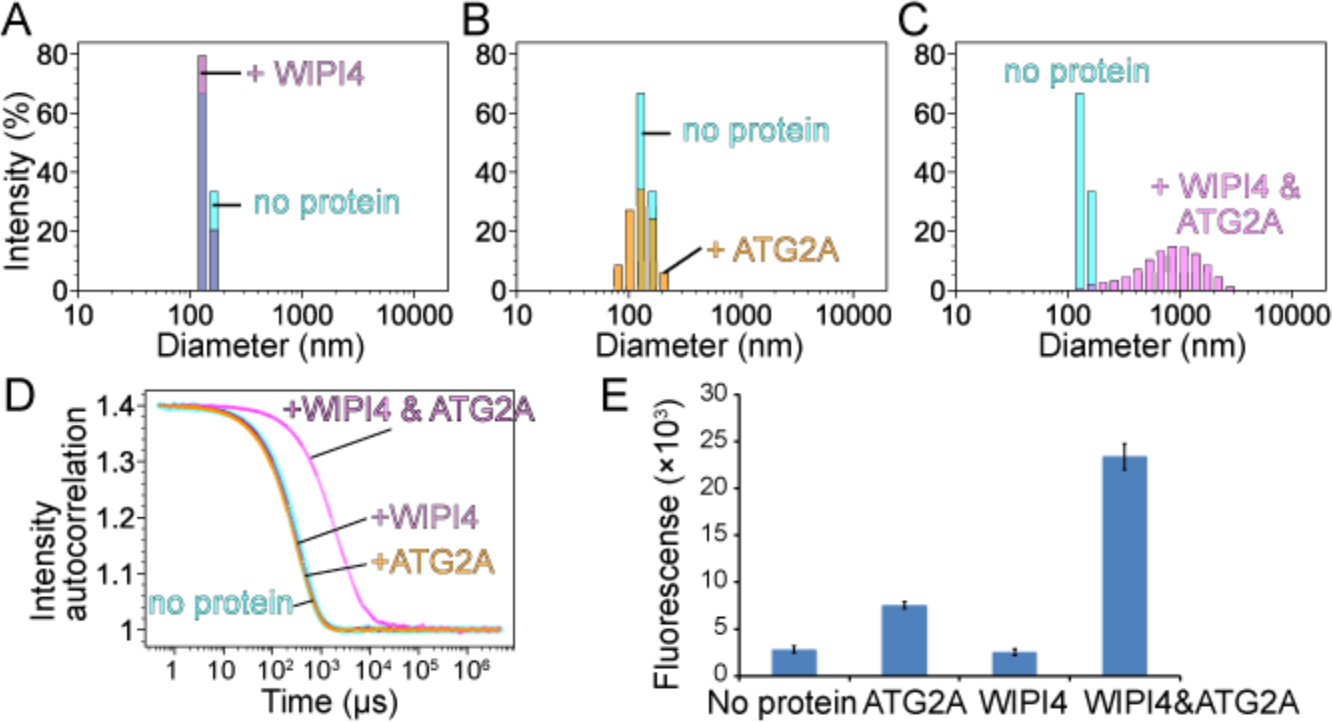
Tethering of PI3P-containing LUVs by the ATG2-WIPI4 complex. (A-C) DLS profiles of LUVs (100 nm) consisting of 75% DOPC, 15% DOPS, 10% PI3P in the absence (A-C: cyan) or the presence of (A) 200 nM WIPI4, (B) 200 nM ATG2A, or (C) both proteins. (D) Auto-scaled autocorrelation functions of the four DLS measurements. (E) Fluorescence-based liposome tethering assay. Higher the fluorescence, more associations between the liposomes with and without PI3P. LUVs consisting of 73.3% DOPC, 15% DOPS, 10% PI3P, 0.2% biotinylated lipids, 1.5% Rhodamine-PE were mixed with LUVs consisting of 73% DOPC, 25% DOPS, 2% DiD in the presence of the indicated proteins. For each experiment, the average of three repeats is shown with the standard deviation.

Our structure of the ATG2A-WIPI4 complex in which WIPI4 is located adjacent to only one of the two membrane-binding sites (the CAD tip) suggests that the ATG2A-WIPI4 complex could asymmetrically bridge two membranes – one containing PI3P to one without PI3P - via WIPI4 and the N tip, respectively. To test the capacity for such heterotypic tethering directly, we performed fluorescence tethering assays with LUVs containing PI3P and biotinylated lipids, and with LUVs lacking both. WIPI4 on its own did not change the fluorescence signal compared to the control (Fig. 5E). Addition of ATG2A showed only a slight increase in signal similar to what was observed using non-PI3P containing LUVs (Fig. 4F), demonstrating that the presence of PI3P does not improve ATG2A’s poor ability to tether LUVs. In stark contrast, the signal increased markedly in the presence of the ATG2A-WIPI4 complex, with rises in fluorescence that were on par with what was observed for the homotypic tethering between non-PI3P-containing SUVs (Fig. 4F). Taken together, these data demonstrate heterotypic membrane tethering by the ATG2A-WIPI4 complex.

### The CLR fragment binds to membranes in an amphipathic α-helical conformation

The observation that the CLR in isolation localizes to LDs (13) raises the possibility that the CLR is a membrane-binding domain, as many proteins localize to LDs through direct interaction with the lipid monolayer surface (40). To further characterize the role of CLR in membrane binding, we generated a CLR fragment as an MBP fusion construct, since MBP was required to maintain the solubility of the CLR fragment. In a liposome flotation assay, MBP-CLR was recoverable in the top fraction only in the presence of liposomes, whereas MBP alone was not detected in the top fraction, demonstrating direct membrane binding by the CLR (Fig. 6A). MBP-CLR did not exhibit any preferences for membrane curvature as it associated with both SUVs and LUVs.

**Fig. 6.**
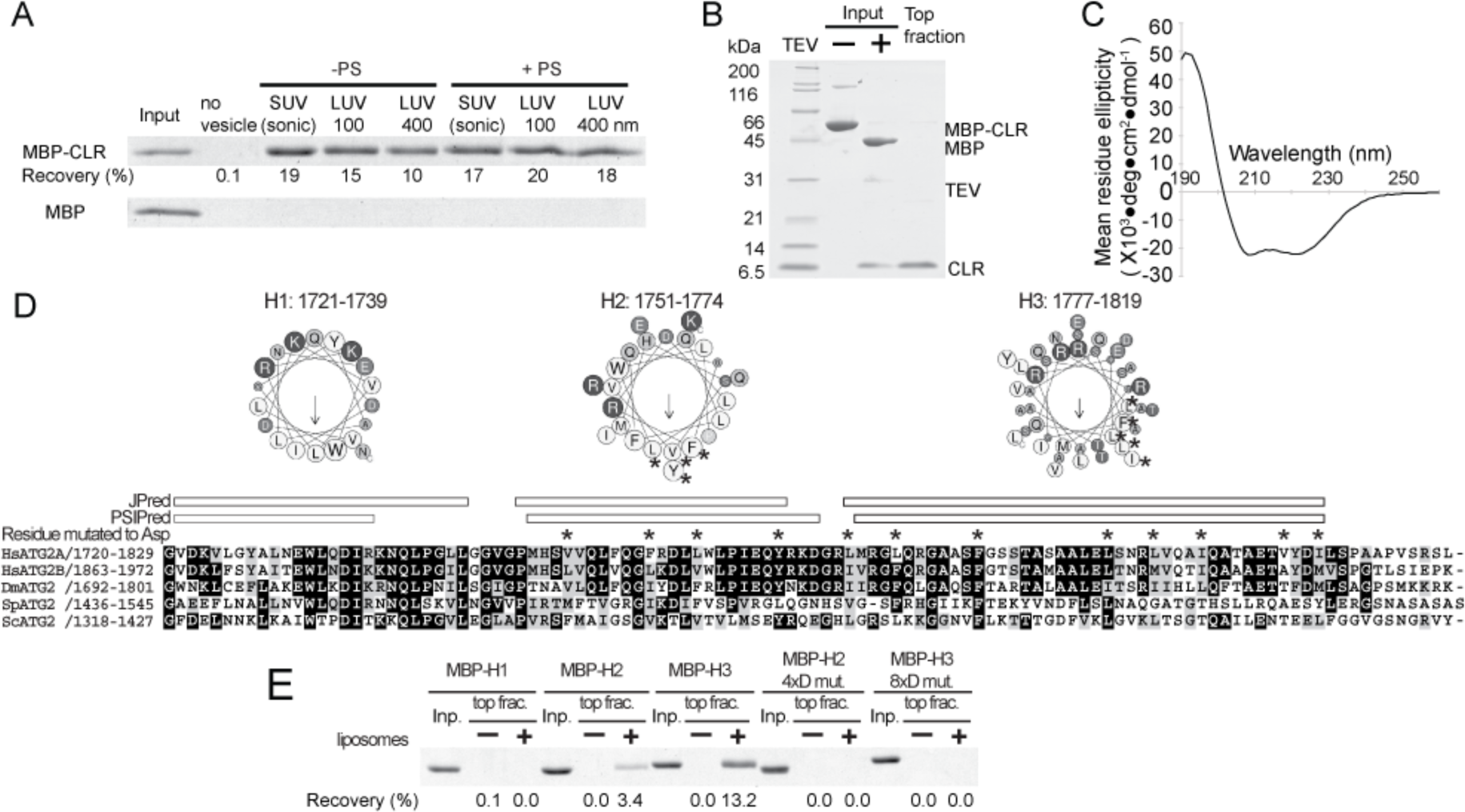
Characterization of the CLR. (A) Liposome flotation assay with 100 nM MBP-fused ATG2A CLR (1723–1819) or MBP alone (control). The liposomes used are indicated as in Fig. 3A. The inputs (4%) and the top layers after the centrifugation (25%) were loaded onto SDS-PAGE. The percentage of MBP-CLR recovered in each of the top fractions was quantified and is shown below the gel image. (B) Preparation of the CLR-SUV complex by liposome flotation. The input containing MBP-CLR and SUVs (75% DOPC and 25% DOPS) was mixed with TEV to cleave off MBP, and the resulting CLR-SUV complex was isolated in the top fraction after centrifugation. (C) The CD spectrum of the CLR-SUV complex. (D) Predicated α-helical regions in the CLR. A multiple sequence alignment (MSA) of the CLR of Homo sapiens (Hs) ATG2A, HsATG2B, Drosophila melanogaster (Dm) ATG2, *Schizosaccharomyces pombe* (Sp) ATG2, and *Sc*ATG2 were generated by clustalw. Secondary structure predictions were obtained using JPred (57) and PSIPred (58) servers. The three fragments generated for flotation assays (H1:1721-1739, H2:1751-1774, H3:1777-1819) are shown as a cartwheel drawing generated by Heliquest server (59). The asterisks shown above the MSA and in the cartwheel drawings indicate the residue mutated to aspartic acid. (E) Liposome flotation assay with 1 µM MBP-fused CLR fragments and LUVs prepared by extruding a lipid mixture consisting of 74% POPC, 25% POPS, and 1% DiD through 100 nm membrane.

Because LD-localized proteins often interact with the LD membrane through their amphipathic α-helices (40), we sought to examine the secondary structure of the CLR. We purified a CLR-SUV complex by removing the MBP tag from the SUV bound MBP-CLR by proteolytic cleavage, followed by liposome flotation (Fig. 6B). Circular dichroism (CD) spectrum of the purified CLR-SUV complex shows a profile typical of α-helix, with local minima at 208 and 220 nm (Fig. 6C). The helical content predicted from the CD spectrum is 61%. A secondary structure prediction suggests that the CLR may contain three α-helical regions, and helical wheel drawings of these regions show that all three (referred to as H1, H2, and H3) may form amphipathic α-helices (Fig. 6D). To determine which region is responsible for membrane binding, we generated each region as an MBP fusion and tested their membrane binding ability. The results of the flotation assay revealed that H2 and H3, but not H1, bind to LUVs consisting of 1-palmitoyl-2-oleoyl-sn-glycero-3-phosphocholine (POPC) and 1-palmitoyl-2-oleoyl-sn-glycero-3-phospho-L-serine (POPS). H3 appears to have a higher affinity for membranes than H2, based on its higher recovery. We then replaced four and eight residues in the hydrophobic side of H2 and H3, respectively, with aspartic acids. These mutations abolished the membrane binding capability of each fragment (Fig. 6E), supporting the likelihood that H2 and H3 bind to membranes via an amphipathic α-helix.

### The CLR is not responsible for membrane tethering by ATG2A

To determine whether the membrane-binding property of the CLR plays a role in membrane tethering, we incorporated all of the mutations described above (a total 12 mutations to aspartic acid) into the full-length ATG2A protein and characterized this mutant (ATG2A^12xD^). We tested membrane binding by flotation assay using liposomes containing PO lipids (POPC and POPS), which were prepared by sonication or extrusion with 30 or 100 nm filter to eliminate potential artifacts (non-specific binding) caused by DO lipids and sonication. ATG2A bound to all types of the liposomes tested but exhibited higher affinity for smaller liposomes (Fig. 7A). These results with DO lipids are consistent with those observed for wild-type ATG2A. The higher preference for sonicated liposomes over the 30 nm liposomes suggests that ATG2A binds to membrane surfaces with local defects rather than sensing overall membrane curvature, as sonication introduces surface defects (37). Our results with ATG2A^12xD^ were very similar to those with the wild-type, indicating these mutated residues are not essential for membrane binding. We then performed membrane tethering assays with ATG2A^12xD^ and found that these mutations do not affect the tethering activity either. The fact that ATG2A^12xD^ clusters SUVs and also mediates the PI3P- and WIPI4-dependent homotypic and heterotypic tethering as efficiently as the wild-type (Fig. 7B–7G) leads us to conclude that the CLR is not involved in the membrane tethering.

**Fig. 7.**
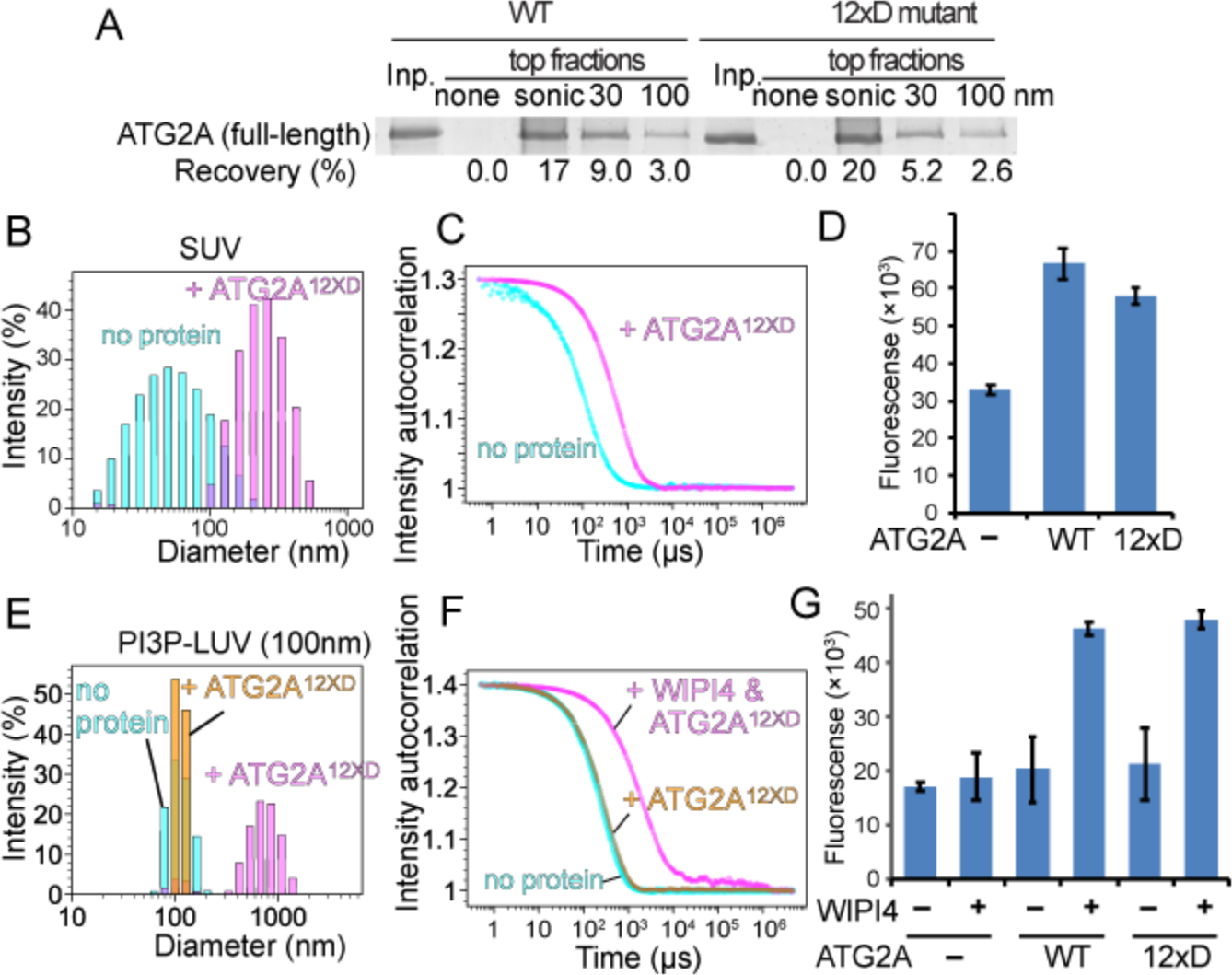
The CLR is not involved in membrane tethering. (A) Flotation assay with ATG2A^12xD^ mutant and liposomes composed of 74% POPC, 25% POPS, and 1% DiD. The liposomes were prepared by sonication or extrusion using 30 or 100 nm filters. The result of SDS-PAGE is shown as in Figs. 3A and 6A. (B, C) DLS homotypic membrane tethering assay with ATG2A^12xD^. The DLS profiles (B) and the autocorrelation functions (C) of the 75%DOPC/25%DOPS SUVs in the absence and presence of ATG2A^12xD^ are shown. (D) Fluorescence homotypic membrane assay with ATG2A wild-type and ATG2A^12xD^ as performed and presented as in Fig. 4F. (E, F) DLS homotypic membrane tethering assay with PI3P-incorporated LUVs (100 nm). The DLS profiles (E) and the autocorrelation functions (F) are shown. (G) Fluorescence heterotypic tethering assay with ATG2A^12xD^ performed in the same manner as shown in Fig. 5E. The experiments with the wild-type ATG2A are repeats of the data shown in Fig. 5E but were performed at the same time as the mutant. Although the fluorescence values are different from those in Fig. 5E, the results from both experiments with the wild-type are consistent with each other.

## Discussion

Recent related studies by Zheng *et al.* on rat ATG2B-WIPI4 complex (41) described the overall shape of the ATG2B-WIPI4 complex by negative stain EM, as well as PI3P-independent membrane binding by ATG2B. They also identified the WIPI4-binding site of ATG2B by a combination of CXL-MS and mutagenesis. Our findings are in agreement with this published work, and their mutational studies reinforce our identification of the WIPI4-binding site on ATG2A. However, here we structurally and biochemically demonstrate that ATG2A is capable of tethering membranes, which provides valuable new insights into autophagosome biogenesis. Gómez-Sánchez et al. also recently characterized the *Sc*ATG2 protein and its interaction with ATG18 and *Sc*ATG9 (42). Their discovery that *Sc*ATG2 binds to membranes by recognizing surface defects is in agreement with our observation that ATG2A binds most strongly with sonicated liposomes. Additionally, their conclusion that *Sc*ATG2 is a mediator of ER-phagophore association complements our structural and biochemical data that demonstrate membrane tethering by the ATG2A-WIPI4 complex.

The membrane organization of the ER-phagophore junction, a site that intricately tied to the omegasome, is poorly understood due to its highly complex and dynamic nature. Thus, predicting the precise location of ATG2A within this junction is a challenge. In Figure 8, we illustrate possible pairings of membranes that may be tethered by the ATG2A-WIPI4 complex at the ER-phagophore junction. The PI3P-enriched omegasome has been suggested to be a cluster of tubular membranes with a diameter of ~30 nm (9), which is similar to the diameters of the SUVs used in this study (Fig. 8A). Our 2D averages of the ATG2A-WIPI4-SUV complex show that the CAD tip and WIPI4 can simultaneously contact the same membrane (Fig. 3B). Thus, it is most logical to assume that the ATG2A-WIPI4 complex associates directly with the omegasome through the CAD tip, as well as through WIPI4 (Fig. 8B). With the CAD tip attached to the omegasome, the N tip could bind to either the ER or the phagophore edge, resulting in a tethering of the omegasome to the ER and/or the phagophore. Alternatively, the N tip may bind to membrane vesicles, such as ATG9 vesicles or COPII vesicles, since ATG2 has been reported to interact with ATG9 and SEC23 (a component of COPII vesicle) (18, 21, 42). These vesicles have been proposed to transform into early phagophores and may also serve as membrane sources for phagophore expansion (6, 43–47). Therefore, tethering of these membrane vesicles by ATG2 to the omegasome would be consistent with the requirement of ATG2 for phagophore expansion (13, 18, 19, 28).

**Fig. 8.**
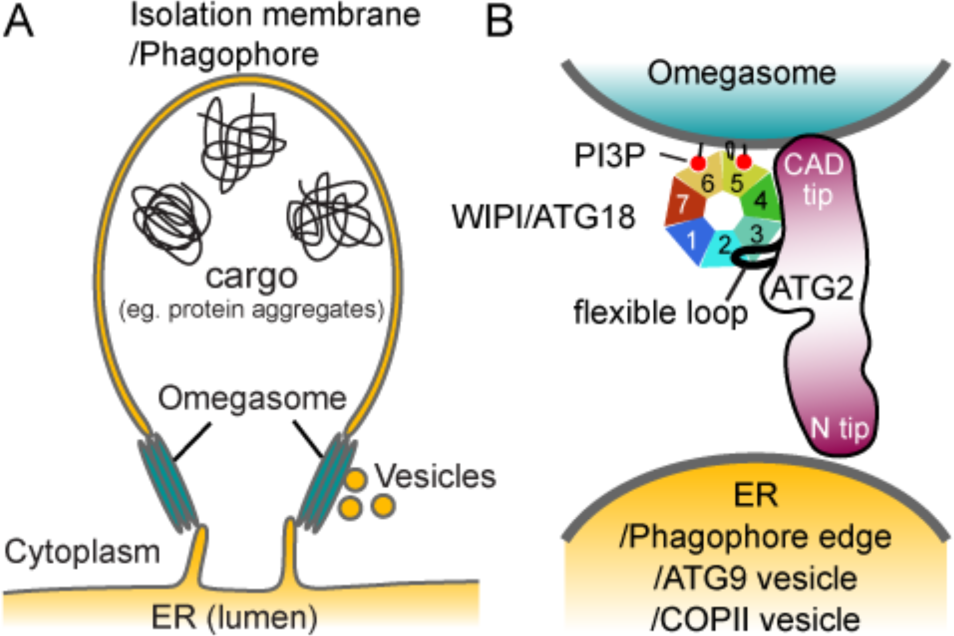
Proposed models of the ER-phagophore/isolation membrane association mediated by the ATG2-WIPI/ATG18 complex. (A) Illustration of the ER-phagophore junction based on current knowledge from cell biological studies. Each gray line represents a lipid bilayer. (B) Structural model of the ATG2-WIPI/ATG18 complex tethering the omegasome to its neighboring membranes (ER, phagophore edge, ATG9 vesicle or COPII vesicle). The blackish red color of ATG2 represents conserved regions as similarly used in Fig. 2A. The WIPI/ATG18 binding region of ATG2 is represented as a black line emanating from the middle region of ATG2 to indicate the flexible attachment of WIPI/ATG18.

Although the precise manner by which ATG2 interacts with this multitude of vesicular components is largely unknown, Gómez-Sánchez et al. report that a short region of *Sc*ATG2 mediates ATG9 interaction (42). This region, located in the reported APT1 domain (23), is partially conserved in ATG2A and starts at the residue 1589 (Figs. 1A and 2A). The closest MBP insertion site amongst our MBP-tagged constructs was at the position 1503-1504, and this MBP was observed adjacent to WIPI4 and the CAD tip in the 2D class averages (Fig. 2B). It is difficult to assume how far this ATG9-binding region is located from the CAD tip, as the 85 residues between the MBP insertion site and the start of the ATG9-binding region could span a long distance if it is flexible. If the ATG9-binding site is located closer to the CAD tip, the CAD tip, ATG18/WIPI, and ATG9 could all be localized to the omegasome. Indeed, under conventional fluorescence microscopy, ATG9 co-localizes with ATG2 and ATG18/WIPI at the ER-phagophore junction (18, 19). However, according to the super-resolution fluorescence microscopy data, the ATG9 compartment and the omegasome are independent units located in the close proximity (6). If the ATG9-binding site is rather closer to the N tip, the ATG2-ATG9 interaction could help stabilize the omegasome-ATG9 compartment association. In either case, there is currently insufficient information to produce a meaningful and complete structural model describing how ATG2, ATG9, and ATG18/WIPI are involved in membrane tethering at the ER-phagophore junction. Higher resolution structural information of these protein complexes and their interactions with membranes, along with more precise localization of these proteins at the ER-phagophore junction will be required to elucidate such a model (48).

Our negative stain EM analysis of the SUV-bound ATG2A-WIPI4 complex suggests that the primary regions in or adjacent to the N-terminus and ATG2_CAD are responsible for membrane binding and tethering (Fig. 3B). Demonstrating membrane interaction of each tip using isolated fragments is particularly challenging, as these fragments are exceptionally recalcitrant to purification. Similarly, obtaining ATG2A proteins containing deletions at these regions has also proven extremely difficult. Such issues have prevented further investigation into the role that these regions play in the membrane tethering process. On a related note, we also attempted to characterize two other previously studied ATG2A constructs having a deletion of the CLR or ATG_C (13, 28), but again failed to obtain sufficient amounts of protein to perform *in vitro* assays. Although the challenges that have been met during the preparation of these truncated proteins may be attributed to sub-optimal constructs or expression and purification conditions, we cannot rule out the possibility that the regions targeted for deletion are integral for the structural stability of the protein. Therefore, site-directed mutagenesis, rather than truncations, would serve as a better strategy for probing the molecular mechanics of membrane tethering, but such studies would require an accurate atomic model of the complexes. While we are unable to provide direct demonstrations of membrane interaction by each tip in isolation, the following evidence supports our conclusions:

First, the N-terminus of VPS13 has been shown to interact with membranes containing negatively charged lipids (49). Thus, the N-terminus of ATG2, which shares sequence similarity with VPS13’s N-terminus, could also be a membrane-interacting domain. Second, the APT1 domain located between ATG2_CAD and the CLR in *Sc*ATG2 has been shown to interact with PI3P-containing membranes (23). Although recent reports (41, 42) and the data we presented here show that ATG2(A/B) bind membranes irrespective of the presence of PI3P, it is still possible that the same region in human ATG2A is responsible for membrane interaction. In that case, the membrane interacting residues in the CAD tip may be the residues in this region, rather than in ATG2_CAD.

In this work, we focused on characterizing yet another conserved region, the CLR. Our data show that the CLR fragment binds to membranes through its two amphipathic α-helices, which is consistent with the LD localization of the CLR fragment (13). However, the mutations in this region that disrupt membrane interaction of the fragment did not affect the membrane tethering activity of the full-length protein (Fig. 7). Recently, a similar set of mutations has been shown to reduce cellular autophagic activity (28). We confirmed that this mutant protein referred to as AH-E (28) is also active *in vitro* membrane tethering (Fig. S5). Thus, the CLR is likely to possess another role essential for phagophore expansion. The membrane tethering activity of ATG2A described in our work cannot alone explain how the phagophore expansion, a process that must involve either lipid transfer or new lipid synthesis, would occur. We speculate that the biochemical function of the CLR may hold the key to this long-standing mystery in autophagosome biogenesis.

Some ATG factors, such as the ATG1 kinase complex, the ATG12-ATG5-ATG16 complex, and ATG8-phosphatidylethanolamine (PE) conjugate, have been reported to mediate membrane tethering *in vitro* (39, 50, 51). While these factors are distinct in molecular organization, they all use protein self-oligomerization to tether neighboring membranes: The ATG1 and ATG12-ATG5-ATG16 complexes both self-dimerize such that two molecules of their membrane-interacting subunit (ATG1 and ATG5, respectively) can associate independently with two separate membranes to tether two vesicles (39, 50, 52). ATG8 is associated with a membrane via its covalent linkage to a PE molecule in the membrane, and multimerization of ATG8 molecules on separate vesicles leads to clustering of the vesicles (51). In contrast, ATG2A has a capacity of bridging two membranes without the requirement for self-oligomerization. ATG2A is rather similar to multi-subunit tethering complexes, such as the Dsl1, HOPS, COG, and TRAPPIII complexes, all of which tether two membranes through the tips of their elongated shapes (53). The roles of the variety of autophagic membrane tethers in autophagosome biogenesis must be different from one other, as they function at different steps. In the earliest step of biogenesis, the ATG1 kinase complex multimerizes ATG9 vesicles, triggering nucleation of a phagophore. The membrane tethering by the ATG12-ATG5-ATG16 complex, which is the E3-ligase like factor for ATG8-PE conjugation, as well as tethering by ATG8-PE, have also been suggested to occur during the nucleation step (44, 50, 51). There is also evidence that ATG8-PE is involved in the final step of phagophore membrane closure (54, 55). ATG2A, which is required for phagophore expansion during the intermediate steps of the autophagosome formation, may be an important collaborator of these other membrane tethers, and begs future study.

## Accession codes

The EM density maps of the ATG2A and ATG2A-WIPI4 complex are deposited to the EMDB with accession codes, 8900 and 8899, respectively.

## Acknowledgments

We thank Drs. Imre Berger and Maria Martinez-Yamout for sharing the multibac system and cDNA of GB1, respectively, Drs. Elizabeth M. Wilson-Kubalek and Zoltan Metlagel for assistance in preliminary EM studies, Drs. Peter E. Wright and Qinghai Zhang for access to CD and DLS instruments, respectively. This research was supported by the grants from the National Institute of Health (GM092740 to T.O. and DP2EB020402 to G.C.L.) and by the European Research Council (ERC) (AdvG grant 670821: Proteomics4D) to R.A. G.C.L. is supported as a Searle Scholar and a Pew Scholar.

## Materials and Methods

### Protein expression and purification

The DNA sequence coding human WIPI4 was cloned into pACEBac1 vector (60) with a PreScission protease-cleavable N-terminal histidine tag. Baculoviruses were generated from the resulting vector in *Spodoptera frugiperda* 9 (Sf9) cells using multibac system (60). Sf9 cells infected with the baculoviruses were grown at 27˚C, harvested 48 hours post-infection, resuspended in lysis buffer 20 mM HEPES pH7.5, 150 mM NaCl, 0.5 mM TCEP, and 1 mM PMSF, and lysed using a Dounce homogenizer. The lysate was clarified by centrifugation at 40,000 × *g*, and the protein in the supernatant was purified by nickel affinity chromatography, followed by anion exchange chromatography on a Source 15Q column (GE Healthcare) and size exclusion chromatography on a Superdex 200 column (GE Healthcare) equilibrated with 10 mM HEPES pH7.5, 300 mM NaCl, and 0.5 mM TCEP. The eluted protein was concentrated, flash-frozen in liquid nitrogen, and stored in a −80˚C freezer until use. WIPI4 without the histidine tag was generated by cleaving the tag between the nickel affinity and the anion exchange chromatography steps. ATG18 cDNA sequence was cloned into pFastBacHT (Life Technologies) with a TEV protease-cleavable N-terminal tag. His-ATG18 protein was expressed in Sf9 cells using baculoviruses generated with this vector as described above and purified by nickel affinity, Source 15Q anion exchange, Phenyl HP hydrophobic Sepharose (GE Healthcare) and Superdex 200 size exclusion chromatographic methods, followed by removal of the histidine tag by TEV protease. Upon digestion, TEV protease was removed by second nickel affinity chromatography (TEV is tagged with the histidine tag). The ATG18 protein in the flow-through of the nickel beads was frozen and stored as described above.

Human ATG2A and *Sc*ATG2 cDNA sequences were cloned into pACEBac1 vector with a TEV-cleavable glutathione-S-transferase (GST) and a PreScission-cleavable Twin-StrepII (TS) tags at the N- and C-terminus, respectively. Baculoviruses were generated using the resulting vectors and the proteins were expressed in Sf9 cells as described above. Cells were harvested, resuspended in lysis buffer 20 mM HEPES pH7.5, 150 mM NaCl, 0.5 mM TCEP, and 1 mM PMSF, and lysed by adding 2% Triton X-100, followed by 1 hour stir-mixing at 4˚C. The lysate was clarified by centrifugation at 40,000 × *g*. The supernatant was diluted to reduce the concentration of Triton X-100 below 1% and to contain total 500 mM NaCl and then mixed with Glutathione Sepharose beads (GE Healthcare) in a batch mode. After a 4-hour incubation at 4˚C, the beads were collected by centrifugation and transferred into a gravity column. The beads were washed with buffer containing 20 mM Tris-HCl pH 8.0, 500 mM NaCl, 0.5 mM TCEP, 25% glycerol, and 0.1% Triton X-100, then TEV protease was added to the beads for removal of the GST tag. The cleaved protein was eluted in the same buffer, flash-frozen in liquid nitrogen and stored at −80˚C until further use. For negative stain EM analysis of ATG2A alone, the protein solution was thawed and loaded onto Strep-Tactin Superflow affinity chromatography beads (IBA). The beads were washed extensively with 100 mM Tris-HCl pH 8.0, 150 mM NaCl, 0.5 mM TCEP to remove Triton X-100, followed by elution with the same buffer containing additional 5 mM desthiobiotin. For the negative stain EM analysis of the ATG2A-WIPI4 and *Sc*ATG2-ATG18 complexes, the Strep-Tactin Superflow beads loaded with hATG2A or *Sc*ATG2 were mixed with an excess amount of WIPI4 or ATG18. After a brief incubation, the beads were washed and eluted as described above. The proteins eluted from the Strep-Tactin Superflow were used immediately for preparations of negative stain EM grids. For membrane binding and tethering assays, the elution from Glutathione Sepharose beads was thawed and loaded onto Strep-Tactin XT Superflow beads (IBA). The beads were washed, and protein was eluted similarly as described above except that 50 mM biotin instead of desthiobiotin was used for the elution. PreScission protease was added to the elution for removal of the C-terminal TS tag. The cleaved tag and biotin were removed by dialysis with three-time buffer exchanges. The purified ATG2A protein was used immediately for biochemical assays. The ATG2A^12xD^ mutant, in which all of the aspartic acid mutations in H2 and H3 regions described below were incorporated (total 12 mutations), was expressed and purified from the baculovirus-infected Sf9 cells in the same manner as the wild-type protein.

For the N-terminal MBP labeling, the DNA sequence coding ATG2A was cloned into pFastBacHT vector with a non-protease-cleavable N-terminal MBP tag and a TEV-cleavable C-terminal histidine tag. The MBP-ATG2A-His protein was expressed in insect cells similarly as described above. 10 mM imidazole pH8.0 was added to the clarified cell lysate, which had been prepared as described above, and the lysate was mixed with TALON cobalt affinity beads (Clontech Laboratories). After a one-hour incubation at 4˚C, the beads were collected and washed with 20 mM Tris-HCl pH 7.5, 300 mM NaCl, 10 mM imidazole, 0.5 mM TCEP, 20% glycerol, and 0.05% Triton X-100, followed by elution with the same buffer except containing 150 mM imidazole. The eluate was then loaded onto Amylose affinity beads (New England Biolabs). After a wash with 20 mM Tris-HCl pH 8.0, 500 mM NaCl, 0.5 mM TCEP, 20% glycerol, and 0.1% Triton X-100, an excess amount of WIPI4 was added to the beads, followed by a brief incubation and rewashed with the 20 mM Tris-HCl pH 7.5, 150 mM NaCl, 0.5 mM TCEP, 10% glycerol, and 0.05% Triton X-100. The protein was eluted with the same buffer containing additional 10 mM maltose and used immediately for EM grid preparation.

For the internal MBP labeling, the DNA fragment coding the amino acid sequence Gly-Gly-Ser-Gly-Ala-Ser-MBP sequence-Ser-Gly-Gly-Ser-Gly-Gly was inserted into the positions between 1224 and 1225 (in ATG2_CAD), 1344 and 1345, 1373 and 1374, 1503 and 1504, and 1829 and 1830 (between the CLR and ATG_C) of ATG2A in pACEBac1 vector harboring GST-ATG2A-TS described above. The MBP-inserted proteins were purified similarly as the wild-type by Glutathione Sepharose chromatography, followed by on-beads TEV cleavage for removal of the N-terminal GST tag. Before the TEV addition, the beads were washed with 20 mM Tris-HCl pH 7.5, 1 M NaCl, 20% glycerol, 0.5 mM TCEP, and 0.026% DDM. The cleaved proteins were eluted in this buffer and concentrated, and then injected into Superose 6 size exclusion column equilibrated with the same buffer. The fractions containing MBP-inserted ATG2A were pooled and concentrated. An excess amount of WIPI4 was added and the complex was isolated in Superose 6 size exclusion column equilibrated with 20 mM Tris-HCl pH 7.5, 500 mM NaCl, 0.5 mM TCEP, 10% glycerol, and 0.026% DDM. The proteins eluted from this column were immediately used for the preparation of EM grids.

For the C-terminal MBP labeling, the DNA sequence coding MBP was inserted between PreScission protease recognition sequence and TS tag of the pACEBac1 vector harboring GST-ATG2A-TS fusion. The GST-ATG2A-MBP-TS protein was expressed and purified in the same manner as the wild-type ATG2A. Before the elution from Strep-Tactin XT affinity beads, WIPI4 was added to form the complex. After wash with 20 mM Tris-HCl pH 7.5, 150 mM NaCl, 0.5 mM TCEP, 10% glycerol, and 0.026% DDM, the protein was eluted in the same buffer supplemented with 50 mM biotin and used for EM analysis.

The DNA sequences coding the CLR fragments (CLR: residues 1723-1819; H1: 1721-1739; H2: 1751-1774; H3: 1777-1819) were cloned into pET Duet-1 vector (Novagen) with a TEV-cleavable N-terminal His-MBP tag and the *Mycobacterium xenopi* gyrA intein-chitin binding domain (CBD) fusion unit from TXB1 vector (New England Biolabs). *E. coli* BL21(λDE3) cells were transformed with the resulting vector harboring His-MBP-CLR-intein-CBD under T7 promoter and grown in Terrific broth medium at 37˚C. Protein expression was induced by adding 0.4 mM IPTG when OD_600_ reached ~2.4. Cells were further grown for overnight at 22C˚, harvested, resuspended in 20 mM Tris-HCl pH 8.5, and 500 mM NaCl, and lysed using an Avestin C3 homogenizer. The lysate was clarified by centrifugation at 40,000 × *g*. 0.2 (v/v) % Triton X-100 was added the supernatant, which was mixed with chitin affinity beads (New England Biolabs) in a batch mode for 2 hours at 4˚C. The beads were collected into a gravity column and washed with 20 mM Tris-HCl pH 8.5, 500 mM NaCl, and 0.1% Triton X-100. 40-50 mM DTT was added to the washed beads for intein cleavage and the beads were incubated at room temperature for initial 5 hours, followed by overnight incubation at 4˚C. Subsequently, His-MBP-CLR was eluted in the same buffer, concentrated and injected into a Superdex 200 size exclusion column equilibrated with 20 mM Tris-HCl pH 7.5, 500 mM NaCl, and 0.05% Triton X-100. The fractions containing the proteins were loaded onto Amylose affinity beads (New England Biolabs) and washed with 20 mM Tris-HCl, 150 mM NaCl extensively to remove detergents. The proteins were eluted in the same buffer supplemented with 10 mM maltose and used immediately for membrane binding assays and CD experiments. The MBP-fused H2 and H3 fragments with 4 (V1754D/F1761D/L1765D/Y1772D) and 8 (L1778D/L1782D/F1789D/L1800D/L1804D/I1808D/V1815D/L1819D) aspartic acid mutations were generated in the same manner as the wild-type fragments as described above.

The DNA sequence coding the residues 1358-1404 of ATG2A (the WIPI4-binding fragment) was cloned into pET vector with a His-GB1 N-terminal tag. The fusion protein was expressed in *E. coli* BL21(λDE3) similarly as described above for the CLR fragments and purified by nickel affinity chromatography, followed by size exclusion chromatography on a Superdex 200 column. The purified proteins were concentrated and stored at −80˚C until use.

### Affinity capture binding assay of the ATG2-WIPI4 and ATG2-ATG18 complexes

Strep-Tactin Superflow beads loaded with ATG2A were mixed with an excess amount of His-WIPI4 in buffer containing 20 mM HEPES pH 7.5, 150 mM NaCl, 0.5 mM TCEP, 20% glycerol, and 0.1% Triton X-100. Beads were collected by centrifugation and washed with the same buffer three times. Proteins were eluted with the same buffer containing additional 5 mM desthiobiotin. The contents of the elution were examined by SDS-PAGE analysis (Fig. 1B). The control was carried out using the beads without ATG2A.

### Reconstitution of the ATG2A-WIPI4-SUV complex for negative stain EM analysis

The lipid mixture (75% DOPC/25% DOPS) was dried under nitrogen gas and vacuum dried. The obtained lipid film was dissolved in a buffer 20 mM Tris pH 8.0, 150 mM NaCl, and 50 mM sodium cholate and dialyzed against the same buffer without sodium cholate. This dialysis procedure produced SUVs that were more homogenous than sonication method. The diameter of the SUVs was ~30 nm as determined by EM and DLS. To reconstitute the protein-SUV complex, ATG2A-TS was loaded onto StrepTactin beads. After a wash with 20 mM Tris pH 8.0, 150 mM NaCl, and 0.5 mM TCEP, WIPI4 was added to the beads. After 5 min incubation, the beads were rewashed, and the liposomes were added, followed by another 10 min incubation. The beads were then washed with five column volumes, and the protein was eluted with 5 mM desthiobiotin. The eluted ATG2A-WIPI4-SUV complex was observed by negative stain EM as described below. Mixing the purified ATG2A-WIPI4 complex and the SUVs in solution also yielded a similar sample as determined by the negative stain EM while the data was collected using the sample reconstituted on beads (see below).

### Negative Stain Electron Microscopy

#### Sample preparation

Continuous carbon grids for negative stain analyses were prepared by evaporating carbon onto nitrocellulose-coated 400 mesh Cu-Rh maxtaform grids (Electron Microscopy Sciences). For preparing each stained grid, a freshly glow-discharged grid was placed inversely on a 4 µL droplet of purified protein sample, which was sitting on a sheet of Parafilm, and incubated for 2 minutes. Most of the sample was wicked off with a filter paper (Whatman No. 1), and grids were immediately placed onto the surface of a 4 µL droplet of 2% (w/v) uranyl formate solution. After 5 seconds incubation, the grid was removed from the droplet, and excess stain was wicked off from the grid using filter paper. This staining step was repeated two more times. In the last staining step, the stain incubation time was extended to 30 seconds to thoroughly embed the protein sample in stain. Finally, after wicking off bulk stain, the grid was completely air-dried on a piece of Parafilm. Grids for the protein-liposome complex were prepared similarly except that the initial incubation with the protein-liposome sample was extended to 5 minutes and 1% (w/v) uranyl acetate solution was used for staining.

#### Data acquisition

All the electron microscopy data were collected on a Tecnai Spirit (FEI) transmission electron microscope operating at 120keV, using a Tietz F416 CMOS, 4K × 4K camera (TVIPS). Data were acquired using the Leginon automated data acquisition system(61), using a nominal magnification of 52,000 ×, which yielded a pixel size of 2.05 Å at the detector level. Micrographs were collected using a total dose of ~30 electrons/Å^2^, and with a defocus ranging from 1 µm to 1.5 µm. 1,407, 590, 441, 414, 626, 348, 440, 291, 409 and 1,805 micrographs were collected for ATG2A-WIPI4 complex, ATG2A alone, N-terminal-MBP tagged ATG2A-WIPI4 complex, C-terminal MBP-tagged ATG2A-WIPI4 complex, MBP tag at CADS (position 1224) of ATG2A-WIPI4 complex, MBP tag at position 1829 of ATG2A-WIPI4 complex, MBP tag at position 1503 of ATG2A-WIPI4 complex, MBP tag at position 1344 of ATG2A-WIPI4 complex, MBP tag at position 1373 of ATG2A-WIPI4 complex, and ATG2A-WIPI4 complex associated with liposome datasets respectively.

#### Image Processing

The Appion image-processing pipeline (62) was used to process the micrographs for all the datasets. CTFFind3 (63) was used for determining the contrast transfer function (CTF) of each micrograph. Particles were picked from the micrographs using Difference of Gaussians (DoG)-based automated particle picker (64), except the liposome-associated ATG2A-WIPI4 dataset, for which particles were manually picked from micrographs using the Manual Picker program in Appion. The phases of the micrographs were flipped using EMAN (65), and particles were extracted using a box size of 192 × 192 pixels for ATG2A, the ATG2A-WIPI4 complex, and yeast ATG2-ATG18 complex. A box size of 224 × 224 pixels was used for the extraction of all MBP-tagged ATG2A-WIPI4 complexes. For the liposome-bound ATG2A-WIPI4 complex dataset, a 160 × 160 pixel box, which incorporated the entire protein but only a fraction of liposome densities, was used. Particles for all the datasets were binned by a factor of two to speed up image processing. The extracted particle datasets were subjected to a topology-based reference-free 2D classification (66) in Appion, which was used to remove non-particle features and aggregates. The resulting stacks were used for subsequent analyses. To aid visualization of the flexible attachment of WIPI4 to ATG2A, we performed a masked classification that focused on the region surrounding the WIPI4 density using the Maskiton (67) web-based classification server on a pre-aligned stack of particles. The resulting 2D classes were used to create the movie depicting the flexible attachment of WIPI4 to ATG2 (Movie S1).

RELION1.4 (68) was used for 3D analyses of ATG2A and ATG2A-WIPI4 datasets, which contained 37,021 and 75,239 particles respectively. Particle stacks were normalized by eliminating pixels with values above or below 4.5σ of the mean pixel value using the normalization function in RELION, prior to 3D analysis. For 3D reconstruction of the ATG2A-WIPI4 dataset, a cylinder with similar dimensions (220Å in length and 50Å in diameter) as ATG2A was created using the “MO 3” function in the SPIDER (69) image processing program, which was used as an initial volume for a single-class initial 3D classification run in RELION. The resulting 3D class was used as an initial model for further 3D classification. Two rounds of 3D classification were performed to identify the best-resolved subset of particles. This subset was subsequently subjected to 3D auto-refinement by projection matching in RELION. For 3D processing of the ATG2A dataset, the final ATG2A-WIPI4 reconstruction was used as an initial model after 60Å low-pass filtering. This data set was initially subjected to three rounds of 3D classification, and the best-resolved subset of particles was then subjected to 3D auto refinement. Smoothed binary 3D masks were generated from the final refined volumes, extending the mask density by five pixels and applying an eight-pixel falloff. These masks were used to continue the 3D refinements to convergence. The final 3D reconstructions of ATG2A and the ATG2A-WIPI4 complex (Figs.1F and 1G) had an estimated resolution of 29.4 Å and 30.3 Å (Fig. S1) respectively, as assessed using Gold Standard Fourier Shell Correlation with a cutoff of 0.143. 6,386 and 5,317 particles contributed to the final reconstructions of ATG2A and the ATG2A-WIPI4 complex, respectively. 3D maps were visualized and segmented using UCSF Chimera (70). The homology model of human WIPI4 was built from a crystal structure of *Kluyveromyces lactis* Hsv2p (Protein Data Bank code: 4EXV) using the software Modeller (71) and docked into the ATG2A-WIPI4 density using UCSF Chimera.

Focused 2D analysis of the liposome-bound ATG2A-WIPI4 complexes was performed using a similar methodology as described in earlier work (72), with the ISAC (73) 2D classification program. In this process, initial reference-free 2D classification of all the particles resulted in classes depicting the ATG2-WIPI4 complex either tethering two liposomes or attached to a single liposome. Particles belonging to these two types of classes were separated into two sub-stacks, each of which was subjected to reference-free 2D alignment using “sxali2d” program of the SPARX EM data processing package (74). In this initial classification, the liposome electron microscopy density dominated the alignments. Based on the 2D averages of these aligned particles, binary masks were generated to eliminate density corresponding to the liposomes, enabling the 2D ISAC analyses to focus on the signal corresponding to the ATG2-WIPI4 complex. This resulted in class averages where the ATG2A-WIPI4 complexes were better resolved. The translational shifts and rotations determined in the masked analysis were then applied to the corresponding unmasked particles to generate unmasked 2D class averages containing well-resolved ATG2-WIPI4 complexes along with the associated liposomes (Fig. 3B (top row)). The ATG2-WIPI4 3D reconstructions were overlaid on these focused 2D class averages to annotate the different orientations of ATG2A (yellow density) -WIPI4 (green density) relative to the liposome (Fig. 3B (bottom row)).

#### Liposome flotation assay

Lipids (1,2-dioleoyl-*sn*-glycero-3-phosphocholine: DOPC and 1,2-dioleoyl-*sn-*glycero-3-phospho-L-serine: DOPS from Avanti Polar Lipids) and a dye, 1,1’-Dioctadecyl-3,3,3’,3’-tetramethylindodicarbocyanine perchlorate (DiD) (Marker Gene Technologies), in chloroform were mixed at a molar ratio of DOPC/DOPS/DiD=99/0/1 or 74:25:1, or POPC/POPS/DiD=74/25/1, and dried under nitrogen gas. DiD was included to enable visualization of the liposomes. The obtained lipid films were further dried under vacuum for 15-60 minutes, followed by hydration with a buffer containing 20 mM Tris pH 7.5, and 150 mM NaCl. After hydration, the lipids were freeze-thawed seven times using liquid nitrogen and a 42˚C water bath. For SUVs, the lipids were sonicated for ~90 seconds using a probe-type sonicator until the solution becomes translucent. For LUVs, the lipids were extruded through a polycarbonate filter with a pore size of 30, 100 or 400 nm using an Avanti extruder. The flotation was carried out as reported previously (37). In brief, a 300 µL mixture solution of 50 nM ATG2A, 300 µM liposomes and 40% (w/v) Nycodenz (Accurate Chemical) in 20 mM Tris pH 7.5, 150 mM NaCl, and 0.5 mM TCEP was placed at the bottom of a centrifuge tube and a solution 250 µL of 30% Nycodenz in the same buffer was placed above the bottom solution. Finally, 50 µL of the same buffer without Nycodenz was placed on the top. The tubes were centrifuged in an SW55Ti rotor (Beckman Coulter) at 279,982 × *g* (max) for 4 hours at 20˚C. After the centrifugation, the liposomes were visible in the top layer. The top fractions (75 µL) containing all the liposomes were collected, and the contents were examined by SDS-PAGE. The gel was stained with Coomassie Brilliant Blue G-250 and scanned on LI-COR Odyssey infrared imaging system. The bands were quantified using Image Studio 2.0 software.

#### CXL-MS analysis

The CXL-MS analysis was carried out as described previously (33, 34). The ATG2A-WIPI4 complex was prepared at a concentration of 0.5 mg/ml in buffer containing 10 mM HEPES pH 7.5, 500 mM NaCl, 0.5 mM TCEP, 0.026 %(v/v) n-Dodecyl β-D-maltoside (DDM) and 20%(v/v) glycerol. For lysine-specific cross-linking, the pH was adjusted to 8.0 using 1 M HEPES pH 8.5. An equimolar mixture of DSS-d_0_ and DSS-d_12_ (Creative Molecules Inc.) was added to the protein at a concentration of 0.25 mM and incubated at 37˚C for 30 min. For cross-^linking of carboxyl groups and zero-length cross-linking, PDH-d0/d10 and DMTMM (both from^ Sigma-Aldrich) were added to final concentrations of 8.3 and 12 mg/mL, respectively, and the samples were also incubated at 37 °C for 30 min. The cross-linking reactions were stopped by diluting the samples four-fold with DDM-free buffer and removing the excess of cross-linking reagents with the help of Zeba gel filtration spin desalting columns (7K MWCO, Thermo Scientific). This step also helped to remove DDM, which could interfere with further processing steps and MS analysis. The filtrate was dried in a vacuum centrifuge. Samples were reduced, alkylated, and digestion with endoproteinase Lys-C and trypsin according to standard procedures, and purified digests were separated by peptide-level size-exclusion chromatography (Supexdex Peptide 3.2/30, GE). SEC fractions were analyzed by LC-MS/MS on an Easy nLC-1000 HPLC system and an Orbitrap Elite mass spectrometer (both from ThermoFisher Scientific). Data analysis was performed using xQuest (75) against a database containing the sequences of ATG2A, WIPI4 and contaminant proteins (tubulins, heat shock proteins, and keratins).

#### DLS

DLS experiments were carried out with DynaPro Plate Reader II (Wyatt Technology). Solution mixtures containing 200 nM ATG2A and 60 µM (lipid) liposomes (75% DOPC/25% DOPS, or 75% DOPC/15% DOPS/10% 1,2-dioleoyl-*sn*-glycero-3-phospho-(1’-myo-inositol-3’-phosphate) (PI3P)) in buffer 20 mM HEPES pH 7.5, 150 mM NaCl, and 0.5 mM TCEP were incubated for an hour at 25˚C, followed by the DLS measurements with a laser power of 10%. In this condition, the protein alone did not produce measurable scattering, confirming that the DLS signals were from liposomes. Proteinase K (Sigma) was added to the clustered sonicated liposomes at a final concentration of 20 µg/ml, and the DLS was measured after 1 and 2 hours. All data were analyzed in the program DYNAMICS included in the instrument.

#### Fluorescence liposome tethering assay

Fluorescence liposome tethering assays were performed similarly as described previously (39). For the experiments monitoring the tethering by ATG2A alone as shown in Figs. 4 and 7, two kinds of liposomes were prepared. One comprised 73.3% DOPC, 25% DOPS, 0.2% 1,2-distearoyl-*sn*-glycero-3-phosphoethanolamine-N-[biotinyl(polyethylene glycol)-2000] (DSPE-PEG(2000)-Biotin) and 1.5% 1,2-dioleoyl-*sn*-glycero-3-phosphoethanolamine-N-(lissamine rhodamine B sulfonyl) (Rhod-PE), and the other 73% DOPC, 25% DOPS and 2% DiD. SUVs and LUVs were prepared from each lipid mixture by sonication or extrusion. The former and the latter liposomes with the same size were mixed at a concentration of 60 µM (lipid) each in the presence or absence of 100 nM ATG2A. The mixture was incubated at room temperature for 5 min. Streptavidin MagneSphere resin (PROMEGA) was then added to the mixture. After 5 min incubation, the resin was washed five times and resuspended in methanol. DiD fluorescence was normalized against rhodamine fluorescence to correct the difference of the total liposomes absorbed on the beads. For the experiments with PI3P and WIPI4 as shown in Figs. 5 and 7, LUVs with a lipid composition of 73.3% DOPC, 15% DOPS, 10% PI3P, 0.2% DSPE-PEG(2000)-Biotin and 1.5% Rhod-PE were mixed with those of 73% DOPC, 25% DOPS and 2% DiD.

#### CD spectroscopy

SUVs comprised of 75% DOPC and 25% DOPS were prepared by sonication and mixed with MBP-CLR at a molar ratio of protein:lipid=1:50. MBP was cleaved off by TEV protease, and the protein-SUV complex was purified by Nycodenz flotation as described above. The top fraction was dialyzed against the CD buffer consisting of 10 mM potassium phosphate pH 7.4, 150 mM potassium fluoride. After the dialysis, the sample was put into a cuvette with a path length of 1 mm. The final sample contained 6.7 µM CLR. The spectrum was acquired on an Aviv spectrophotometer at 25˚C. A sample containing only SUVs (335 µM lipid) was measured, and the resulting spectrum was used for background subtraction. The secondary structure contents were estimated by the program K2D3 (76).

